# Temporal regulation of cell polarity via the interaction of the Ras GTPase Rsr1 and the scaffold protein Bem1

**DOI:** 10.1101/554105

**Authors:** Kristi E. Miller, Hay-Oak Park

## Abstract

Establishing cell polarity is critical for growth and development of most organisms. The Cdc42 GTPase plays a central role in polarity development in species ranging from yeast to humans. In budding yeast, a specific growth site (i.e. bud site) is selected in the G1 phase, which determines the axis of cell polarization. Rsr1, a Ras GTPase, interacts with Cdc42 and its associated proteins to promote polarized growth at the proper bud site. Yet the mechanism underlying spatial cue-directed cell polarization is not fully understood. Here, we show that Rsr1 associates with Bem1, a scaffold protein, preferentially in its GDP-bound state in early G1. This interaction involves a part of the Bem1 Phox homology (PX) domain, which overlaps with a region previously shown to interact with Exo70, an exocyst component. Furthermore, overexpression of the constitutively GDP-bound Rsr1 interferes with Bem1’s association with Exo70 and inhibits Bem1-dependent Exo70 polarization. We propose that Rsr1 plays a delicate role in coordination of spatial and temporal regulation of polarity establishment via its GTP- and GDP-bound states.

## Introduction

The establishment of polarity and proper positioning of the cell division plane are critical for cell proliferation and development. In the budding yeast *Saccharomyces cerevisiae*, selection of a bud site occurs in a specific pattern depending on cell type and determines the axis of polarized growth. Haploid **a** or α cells bud in an axial pattern, whereas diploid **a**/α cells bud in a bipolar pattern (Freifelder, 1960; Hicks *et al*., 1977; Chant and Pringle, 1995). Selection of a proper bud site depends on cell-type-specific cortical markers and the Rsr1 GTPase module, composed of Rsr1, its GTPase activating protein (GAP) Bud2, and its GDP-GTP exchange factor (GEF) Bud5 (Bender and Pringle, 1989; Chant *et al*., 1991; Chant and Herskowitz, 1991; Park *et al*., 1993). These proteins interact with Cdc42 and its regulators to direct organization of the actin cytoskeleton and septin filaments for polarized growth at the selected site (Bi and Park, 2012).

In the absence of spatial cues, yeast cells can still polarize at a single random site. This spontaneous cell polarization (often referred to as ‘symmetry breaking’) may occur via positive feedback loops involving the actin cytoskeleton or a Cdc42 signaling network that includes Bem1, the Cdc42 GEF Cdc24, and the Cdc42 effector PAK (p21-activated kinase) (Wedlich-Soldner *et al*., 2003; Irazoqui *et al*., 2003; Wedlich-Soldner *et al*., 2004; Goryachev and Pokhilko, 2008; Kozubowski *et al*., 2008). Despite a large number of studies, several aspects of the mechanisms underlying Cdc42 polarization have been under debate (Smith *et al,* 2013; Woods *et al.,* 2015; Rapali *et al.,* 2017). It also remains elusive whether and how these polarity factors may be involved in spatial cue-directed polarization of wild-type (WT) cells, as these studies in spontaneous cell polarization used cells lacking *RSR1*.

Another important aspect that has been unclear is temporal regulation of polarity establishment. The G1 phase in budding yeast is partitioned into two distinct steps, T_1_ and T_2_, by the exit of the transcriptional repressor Whi5 from the nucleus (Di Talia *et al*., 2007). When 50% of Whi5 has exited the nucleus, yeast cells pass through the cell-cycle commitment point, known as ‘Start’ (Doncic *et al*., 2011). We previously found stepwise activation of Cdc42 in relation to these two steps in G1: Cdc42 is activated by Bud3 in early G1 and subsequently by Cdc24 (Kang *et al*., 2014). The Rsr1 GTPase module is necessary for the first step of Cdc42 polarization in haploid cells (Lee *et al*., 2015). Rsr1 also interacts with the polarity proteins: Rsr1-GTP interacts with Cdc24 and Cdc42 (Zheng *et al*., 1995; Park *et al*., 1997; Kozminski *et al*., 2003; Kang *et al*., 2010). Surprisingly, Rsr1 was found to interact with Bem1 preferentially in its GDP-bound state in vitro (Park *et al*., 1997), although the physiological significance of this interaction has been unclear.

Bem1 functions as a signaling hub linking many binding partners that interact with its different domains (Ito *et al*., 2001; Bose *et al*., 2001; Yamaguchi *et al*., 2007; Stahelin *et al*., 2007; Takaku *et al*., 2010; Liu and Novick, 2014). How could these multiple interactions be temporally regulated? Previous studies have suggested that Cdc24 activity is enhanced by Bem1 (Smith *et al*., 2013; Rapali *et al*., 2017), which associates with Cdc24 after Start (Witte *et al*., 2017). Contrary to these studies, a recent report argues that Bem1 and Cdc24 are active and promote Cdc42 polarization before Start (Moran *et al*., 2018). In this study, we sought to resolve these discrepancies and answer the following outstanding questions: When does Bem1 function in spatial cue-directed polarization of WT cells? Does Rsr1 indeed have two active states where the GDP-bound form interacts with Bem1 in vivo? If so, what is the functional significance? To address these questions, we examined Cdc42 polarization and Bem1 localization as well as protein interactions in yeast. Here we report that Bem1 and Rsr1-GDP interact in early G1. We provide evidence that the association of Rsr1-GDP and Bem1 may hinder Bem1-dependent Exo70 localization and delay polarized secretion.

## Results and Discussion

### Rsr1-GDP associates with Bem1 in early G1

To determine whether Rsr1-GDP interacts with Bem1 in vivo, we performed a bimolecular fluorescence complementation (BiFC) assay, which is based on recovery of split fluorescent proteins (Hu *et al*., 2002; Kerppola *et al*., 2009). We expressed Bem1 fused to YN (the N-terminal half of YFP) together with YC (the C-terminal half of YFP) fused to WT Rsr1 and mutant Rsr1^G12V^ and Rsr1^K16N^, which are expected to be in the GTP- and GDP-locked states in vivo, respectively (Ruggieri *et al*., 1992), in haploid cells. YFP fluorescence was observed in cells expressing Bem1-YN and YC-Rsr1^K16N^ (and YC-Rsr1) at the bud neck of large budded cells and at the division site in unbudded cells, whereas little fluorescence was detected in cells expressing Bem1-YN and YC-Rsr1^G12V^ (**Figure 1Aa**). We also observed positive BiFC signals in a control strain expressing Bem1-YN and YC-Cdc42, consistent with previous reports that Bem1 interacts with Cdc42 (Yamaguchi *et al*., 2007; Takaku *et al*., 2010). Similarly, Bem1-YN associated with YC-Rsr1^K16N^ but not with YC-Rsr1^G12V^ in diploid cells (**Figure S5C**). These observations suggest that Rsr1 indeed associates with Bem1 in its GDP-bound state in vivo.

**Figure 1.**
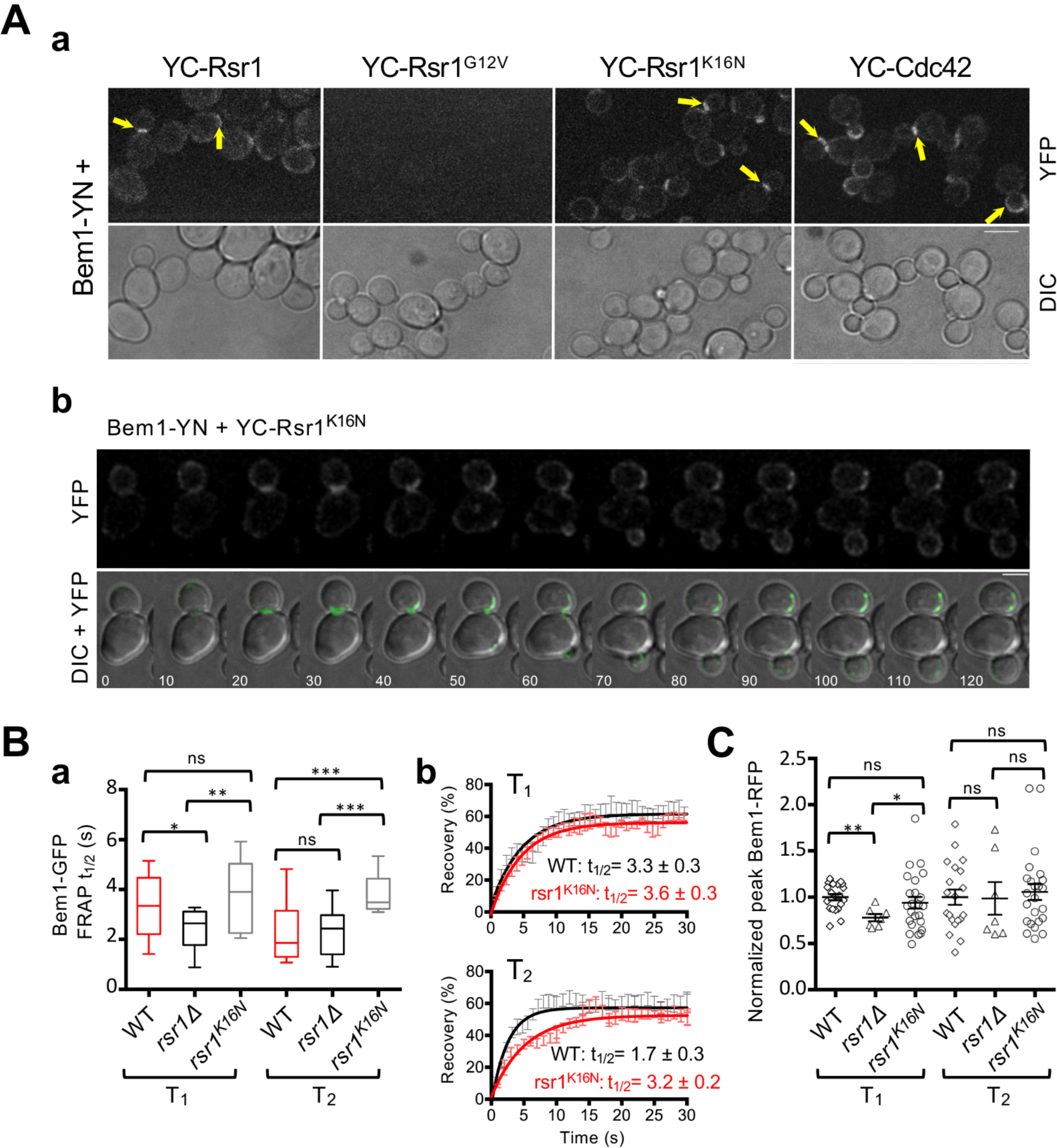
Rsr1-GDP may associate with Bem1 during T_1_. (A) (a) BiFC assays in haploid cells expressing YC-Rsr1, YC-Rsr1^G12V^, YC-Rsr1^K16N^, or VC-Cdc42, along with Bem1-YN. Bar, 5µm. (b) Time-lapse images of cells expressing YC-Rsr1^K16N^ and carrying pRS426-Bem1-YN at 22°C. Numbers indicate time (min) from the first capture. Bar, 3µm. (B) (a) FRAP analysis of Bem1-GFP at the division site during T_1_ (WT, n =11; *rsr1*Δ, n =12; and *rsr1K16N*, n =14) or at the incipient bud site during T_2_ (WT, n =16; *rsr1*Δ, n =13; and *rsr1^K16N^*, n =9). (b) FRAP curves of Bem1-GFP in WT or *rsr1^K16N^* cells during T_1_ or T_2_. (C) Peak Bem1-RFP localized intensity during T_1_ or T_2_ is plotted for individual daughter cells (WT, n =20; *rsr1*Δ, n =7; and *rsr1^K16N^*, n =24).

As expected from static images (**Figure 1Aa**), time-lapse imaging of cells expressing Bem1-YN and YC-Rsr1^K16N^ suggested that Bem1 and Rsr1^K16N^ might interact in late M and early G1 (**Figure 1Ab**). Because a potential caveat of BiFC assays is irreversible association of fusion proteins (Kerppola *et al*., 2009), we also used fluorescence recovery after photobleaching (FRAP) with cells expressing GFP-tagged Bem1 (and Whi5-RFP as a cell cycle marker) to address when in the cell cycle Rsr1-GDP interacts with Bem1. If Bem1 associates with Rsr1 at a specific stage in G1, dynamics of Bem1-GFP might be affected in *rsr1* mutants compared to WT during T_1_ or T_2_. Indeed, we found that Bem1-GFP recovers faster after photobleaching in *rsr1*Δ cells than in WT during T_1_ (i.e., when Whi5 was in the nucleus), while Bem1-GFP dynamics were similar during T_2_ in these cells (**Figure 1Ba**). These data suggest that Rsr1 interacts with Bem1 during T_1_ but not after the T_1_-T_2_ transition. Interestingly, Bem1-GFP exhibited slower dynamics during T_2_ in *rsr1*^*K16N*^ cells than in WT (**Figure 1, Ba & Bb**), suggesting that expression of the constitutively GDP-bound or nucleotide-empty Rsr1 may continue to hold Bem1 longer in G1.

### Bem1 co-localizes with Cdc24 and Cdc42-GTP throughout G1 in haploid and diploid cells

Our results discussed above suggest that Rsr1-GDP is likely to associate with Bem1 in early G1. However, when Bem1 localizes has been unclear in haploid cells. Even in diploid cells, which were often used to investigate Cdc42 polarization during symmetry breaking (i.e., in *rsr1*Δ cells), when Bem1 associates with Cdc24 or polarizes has been under debate (Witte *et al*., 2017; Moran *et al*., 2018). To clarify these discrepancies and gain insight into the timing of Bem1’s function in spatial cue-directed cell polarization, we examined Bem1 localization in WT haploid cells together with Whi5-GFP or other polarity markers, Cdc24-GFP and PBD-RFP (the p21-binding domain fused to tdTomato), a biosensor for Cdc42-GTP (Ozbudak *et al*., 2005; Tong *et al*., 2007; Okada *et al*., 2013). Our analyses focused on daughter cells, which have longer T_1_ length compared to mother cells. Bem1-RFP localized to the division site shortly after Whi5-GFP entered the nucleus and then to the incipient bud site around the T_1_-T_2_ transition, and this localization pattern overlapped with the Cdc42-GTP cluster throughout G1 in WT cells (**Figure S1, a & c**). Co-localization of Bem1-RFP with Cdc24-GFP to the incipient bud site was evident during T_2_ (**Figure S1d**), as the majority of Cdc24 resides in the nucleus during early G1 in haploid cells as previously reported (Toenjes *et al*., 1999; Nern and Arkowitz, 2000; Shimada *et al*., 2000). We observed weak Cdc24-GFP signal around the division site during T_1_ and also detected the Cdc24-Bem1 complex by BiFC assays (see below), suggesting that a minor portion of Cdc24 is able to interact with Bem1 at the division site in T_1_. However, this complex may not be active in T_1_ because Cdc42 activation in early G1 is mostly independent of Cdc24 in WT haploid cells (Kang *et al*., 2014) (see more discussion below).

Close examination of Bem1 localization in haploid *rsr1* mutants indicated that the Bem1-RFP peak intensity was reduced in *rsr1*Δ during T_1_ compared to those in WT and *rsr1*^*K16N*^ cells, whereas it was about the same in these cells during T_2_ (**Figures 1C & S1**). These results suggest that Bem1 localization during T_1_ depends on Rsr1-GDP, consistent with the association of Bem1 and Rsr1-GDP. However, Bem1 localization to the division site is not completely abolished in *rsr1*Δ during T_1_, likely due to the presence of other Bem1-interacting protein(s) during this time window.

We then examined localization of Bem1 and these polarity factors in diploid cells to determine how Rsr1 might affect their polarization. Bem1 localized to the division site and to the distal pole (i.e., the pole distal to the birth scar) of diploid daughter cells during T_1_, whereas it became polarized solely at the distal pole after the T_1_-T_2_ transition (**Figures S2a & S3Bb**), as expected from Cdc42 polarization (Lo, Lee *et al*., 2013). In diploid cells, Cdc24 does not localize to the nucleus in late M and early G1, but few cells had Cdc24 at the distal pole in T_1_ (**Figure S3, Aa & Bc**). While Bem1 and Cdc24 co-localize to the division site in early G1, their co-localization to the distal pole of daughter cells was evident only when the incipient bud site was established after T_1_-T_2_ transition (**Figure S3Ab**). In diploid *rsr1*Δ and *rsr1*^*K16N*^ cells, Bem1 localized to the cell division site but was absent from the distal pole during T_1_. In late G1, however, Bem1 localized to the incipient bud site at the distal pole in these *rsr1* mutant daughter cells (**Figures S2, b & c; S3 Ac**). Therefore, Bem1 and Cdc24 localized to the division site is likely functionally inactive, and their localization to the daughter cell’s distal pole in late G1 is independent of Rsr1. Consistently, diploid *rsr1*Δ cells bud at the distal pole in the first budding event (Michelitch and Chant, 1996). These observations suggest that other protein(s) must bring Bem1 and Cdc24 to the cell tip to promote distal pole budding and that Cdc42 polarization during T_2_ is likely independent of Rsr1 at least in diploid daughter cells.

Previous reports also suggest that Cdc42 polarization during T_1_ depends on Bud3 and Rsr1 but is independent of Cdc24 in haploid cells (Kang *et al*., 2014; Lee *et al*., 2015; Kang *et al*., 2018) and that Bem1 may not function in the polarity complex until after Start (Witte *et al*., 2017) (see below for more discussion). On the contrary, another study (Moran *et al*., 2018) argues that Bem1-mediated positive feedback, which includes Cdc24 brought by Rsr1, functions in prestart Cdc42 polarization. The reason of this discrepancy is not clear, but we speculate that hydroxyurea (HU) treatment of cells used by Moran and colleagues might have led to different observations. Arresting cells in early S phase with HU was found to increase Whi5 concentration (Neurohr *et al*., 2018), and thus HU treatment might have resulted in longer T_1_ in the subsequent cell cycle after release, likely affecting localization of the polarity factors in their study.

### Rsr1-GDP may inhibit Bem1’s function in Cdc42 polarization

As described above, we observed that both Bem1 and Cdc24 localize near the division site even during early G1. Then, why might Bem1 not activate Cdc24 in early G1? We hypothesized that the association of Rsr1-GDP with Bem1 in early G1 might hinder Bem1’s function and thus prevent premature activation of Cdc24 until Start. If this were the case, we predicted that Cdc42 polarization during T_2_ might be delayed in *rsr1*^*K16N*^ cells, which express the GDP-locked Rsr1. To test this idea, we monitored Cdc42-GTP polarization in WT and *rsr1* mutants expressing PBP-RFP and Whi5-GFP. Consistent with previous reports (Okada *et al*., 2013; Atkins *et al*., 2013; Kang *et al*., 2014; Lee *et al*., 2015), the Cdc42-GTP level was minimum at the onset of cytokinesis but started to increase soon after cytokinesis (**Figure 2, Aa & Bc**). From analyses of the PBD-RFP cluster, we estimated the time span from the T_1_-T_2_ transition until the Cdc42-GTP level peaked during T_2_ in daughter cells (**Figure 2Ba**). The Cdc42-GTP level reached a maximum slightly earlier in *rsr1*Δ cells after the T_1_-T_2_ transition but was particularly delayed in *rsr1^K16N^* cells – on average 4 mins later –compared to WT cells (**Figure 2B, b & c**). Similarly, the maximum Cdc42 polarization during T_2_ was established about 6 min later in the diploid *rsr1^K16N^* daughter cells compared to the WT diploid daughters (**Figure S4**). Taken together, these results suggest that the expression of GDP-locked Rsr1 delays Cdc42 polarization in daughter cells, although the peak level of Cdc42-GTP cluster during T_2_ was about the same among these strains.

**Figure 2.**
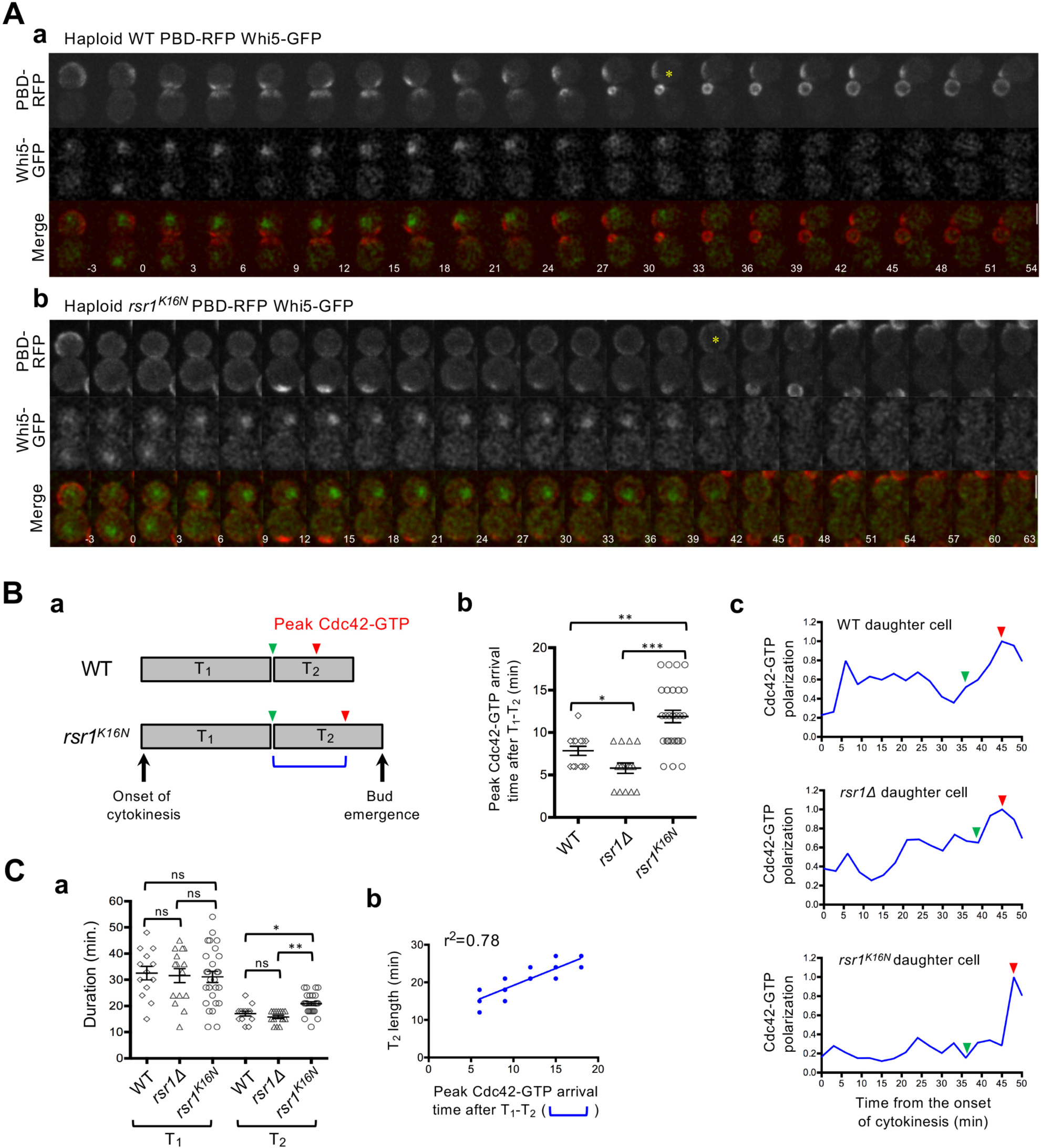
The second phase of Cdc42 polarization is delayed in haploid cells expressing GDP-locked Rsr1. (A) Time-lapse images of PBD-RFP and Whi5-GFP in (a) WT and (b) *rsr1^K16N^* cells at 30°C. Numbers indicate time (min) from the onset of cytokinesis (t=0). Bars, 3 µm. Asterisk marks T_1_-T_2_ transition. (B) (a) Diagram of T_1_ and T_2_ lengths in WT and *rsr1*^*K16N*^ cells. Green and red arrowheads denote T_1_-T_2_ transition and the time at which the Cdc42-GTP cluster reaches a maximum value during T_2_, respectively. The time interval of those two points is marked with a blue bracket and is quantified for individual daughter cells of WT (n=13), *rsr1*Δ (n=15), and *rsr1^K16N^* (n=27) (b). (c) Representative graphs of Cdc42-GTP polarization in daughter cells. Values were normalized to the peak Cdc42-GTP level during T_2_. (C) (a) Length of T_1_ and T_2_ (min) in each daughter cell. (b) Correlation analysis of T_2_ length and the peak Cdc42-GTP arrival time after T_1_-T_2_ transition in *rsr1*^*K16N*^ cells (n=27).

What could be the consequence of delayed Cdc42 polarization? We postulated that delayed Cdc42 polarization in *rsr1^K16N^* cells might result in delayed bud emergence. Indeed, we observed that T_2_ was longer in *rsr1*^*K16N*^ cells compared to WT or *rsr1*Δ cells, while the average T_1_ length was similar among all these strains despite cell-to-cell variations (**Figures 2Ca & S4Ca**). Remarkably, the T_2_ length in individual cells positively correlated with the time when the Cdc42-GTP cluster reached its peak level in T_2_ in both haploid and diploid *rsr1*^*K16N*^ daughter cells (**Figures 2Cb & S4Cb**). Taken together, these observations suggest that the interaction between Rsr1-GDP and Bem1 may interfere with Bem1’s role in mediating Cdc42 polarization.

### Bem1 binds to Rsr1-GDP via its PX domain

As discussed above, Rsr1 may control proper timing of the second phase of Cdc42 polarization by interacting with Bem1. How does Rsr1 regulate Bem1? Bem1 is known to interact with Cdc24 via its PB1 domain (Ito *et al*., 2001) and with Cdc42 and Ste20 through its second SH3 domain and the C-terminal flanking region (aa159-251) (Bose *et al*., 2001; Yamaguchi *et al*., 2007; Takaku *et al*., 2010). The Bem1 PX domain contains a region that interacts with phosphoinositides (Stahelin *et al*., 2007). A region containing both the PX and PB1 domain (aa309-510) of Bem1 has been shown to interact with the exocyst component Exo70 (Liu and Novick, 2014) (**Figure 3Aa**). To gain insight into the mechanism by which Rsr1 regulates Bem1, we first determined by BiFC assays which region of Bem1 binds to Rsr1-GDP. We found that a deletion of the C-terminal half (aa345-408) of the PX domain almost completely abolished the BiFC signal, while a deletion of its N-terminal half (aa281-345) slightly reduced the BiFC signal (**Figure 3**). In contrast, deletions of either the first or second SH3 domain or PB1 domain did not result in an obvious defect in Bem1-YN association with YC-Rsr1^K16N^. Similarly, the K482A mutation in the PB1 domain, which disrupts the interaction between Bem1 and Cdc24 (Ito *et al*., 2001), did not affect the Bem1-Rsr1 interaction (**Figure S5**). These results thus indicate that the C-terminal half of the Bem1 PX domain is required for interaction with Rsr1^K16N^.

**Figure 3.**
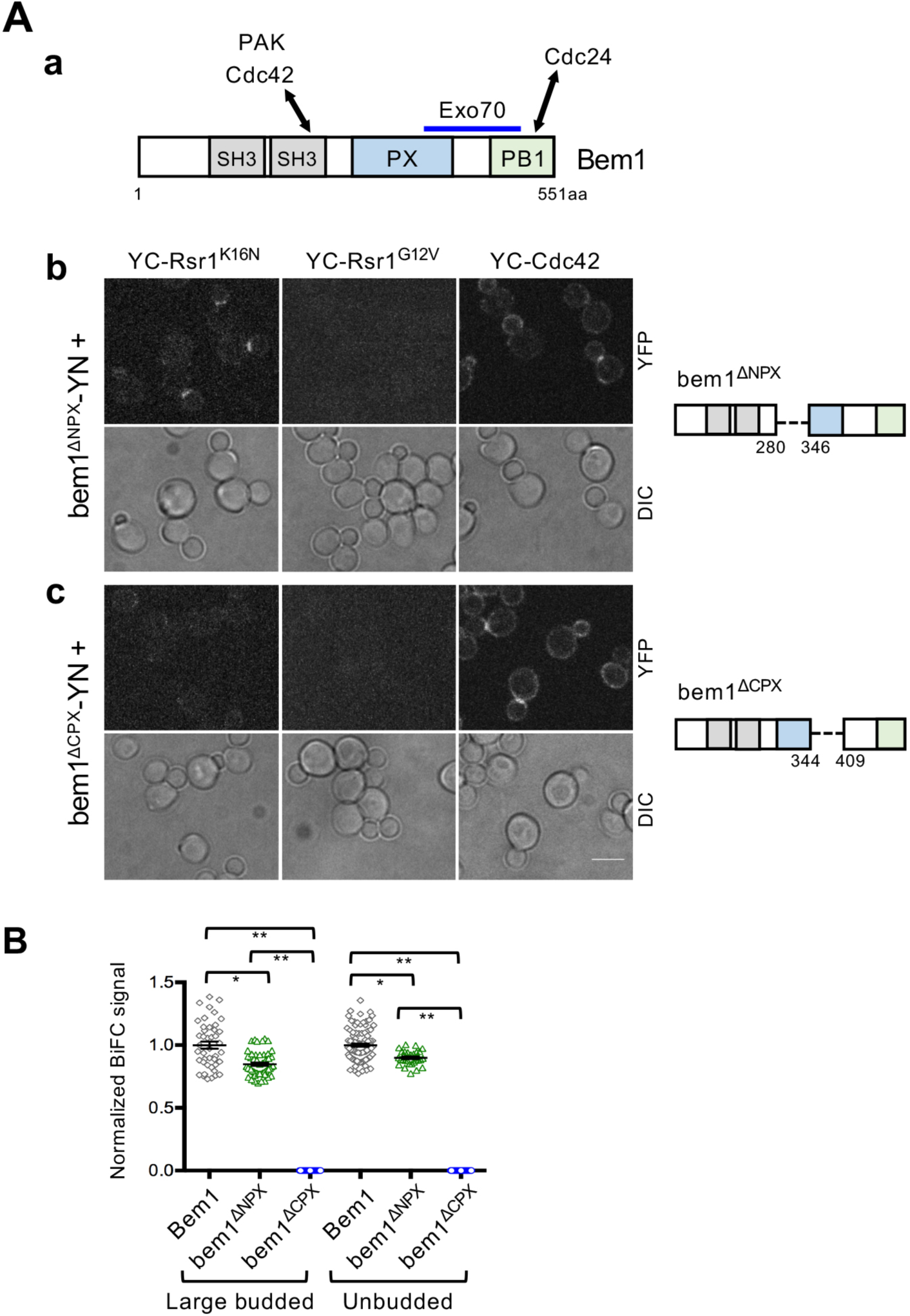
Bem1interacts with Rsr1-GDP likely via its PX domain. (A) (a) Diagram of Bem1 protein domains and known interactions (see text). (b) BiFC assays in haploid cells expressing YC-Rsr1^K16N^, YC-Rsr1^G12V^, or YC-Cdc42 along with (b) bem1^ΔNPX^-YN or (c) bem1^ΔCPX^-YN. See a scheme on the right for each deletion. Bar, 5µm. (B) Quantification of BiFC signals in cells expressing YC-Rsr1^K16N^ and Bem1-YN (large-budded, n=43; unbudded, n=94), bem1^ΔNPX^-YN (large-budded, n=51; unbudded, n=32), or bem1^ΔCPX^-YN (large-budded, n=63; unbudded, n=100).

### Rsr1-GDP may inhibit Bem1’s role in Exo70 polarization

Interestingly, the C-terminal PX domain of Bem1, which is necessary for interaction with Rsr1, overlaps with the region that mediates actin-independent localization of Exo70 (Liu and Novick, 2014). Thus, we hypothesized that Rsr1-GDP associates with Bem1 and inhibits it from promoting Exo70 localization. To test this idea, we examined how overexpression of Rsr1^K16N^ affects Exo70 polarization in cells transiently inhibited for actin polymerization to block actin-dependent delivery of Exo70. We imaged cells expressing Exo70-RFP (and Whi5-GFP) carrying a multicopy Rsr1^K16N^ plasmid or a vector control after treatment with Latrunculin A (LatA), an actin assembly inhibitor (Ayscough *et al*., 1997). Exo70 polarization was not affected by overexpression of Rsr1K16N in cells with a small bud or unbudded cells in T_1_. In contrast, overexpression of Rsr1^K16N^ did cause decreased polarized localization of Exo70 in mock-treated (DMSO) unbudded cells in T_2_, and this decrease was even more pronounced in LatA-treated cells in T_2_ (**Figure 4A**).

**Figure 4.**
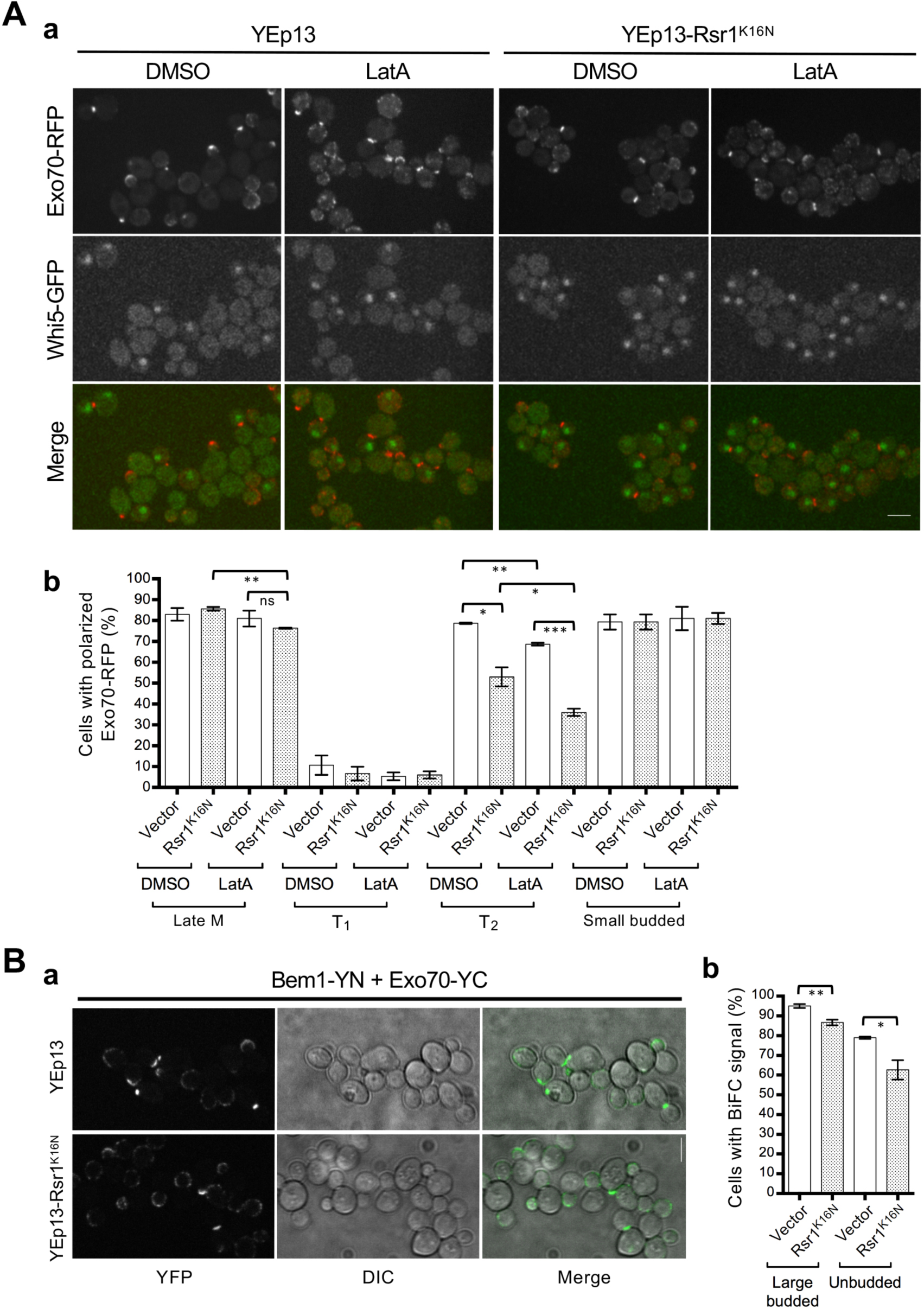
Rsr1-GDP may hinder Bem1-dependent Exo70 polarization. (A) (a) Images of LatA or DMSO treated WT haploid cells expressing Exo70-RFP and Whi5-GFP and carrying each plasmid as marked. Bar 5µm. (b) The percentage of cells with polarized Exo70-RFP (n=100-270 for each sample per experiment). (B) (a) BiFC assays in the haploid *BEM1-YN EXO70-YC* strain carrying YEp13 or YEp13-Rsr1^K16N^. Bar, 5µm. (b) Large-budded or unbudded cells with BiFC signal was quantified from three separate experiments (n=50-130 for each sample per experiment).

In a second approach, we used BiFC assays to determine how overexpression of the GDP-locked Rsr1 affects the Bem1-Exo70 interaction. As expected from a previous study (Liu and Novick, 2014), we observed strong BiFC signals in cells expressing Bem1-YN and Exo70-YC. When these cells were examined after transforming with YEp13-Rsr1^K16N^ or vector control, we found that the number of large budded or unbudded cells with positive BiFC signals decreased when Rsr1^K16N^ was overexpressed (**Figure 4B**), indicating that GDP-locked Rsr1 interferes with the Bem1 and Exo70 interaction. Collectively, these results suggest that Bem1-dependent Exo70 localization is diminished by overexpression of the GDP-locked Rsr1, supporting our hypothesis that Rsr1-GDP inhibits Bem1’s function in promoting Exo70 polarization.

### Rsr1-GDP may not interfere with the Bem1-Cdc24 interaction

Since we observed a delay in Cdc42 polarization in *rsr1^K16N^* cells (see above), we considered the possibility that the interaction between Bem1 and Rsr1-GDP may limit the number of free Bem1 protein in the cell, hindering the Bem1-Cdc24 association. To test this idea, we determined by BiFC assays how Bem1-Cdc24 association was affected by overexpression of *rsr1^K16N^*. While Bem1-YN formed a bimolecular fluorescent complex with Cdc24-YC, as expected, we found that overexpression of Rsr1^K16N^ did not have any obvious effect on the BiFC signals (**Figure 5A**).

**Figure 5.**
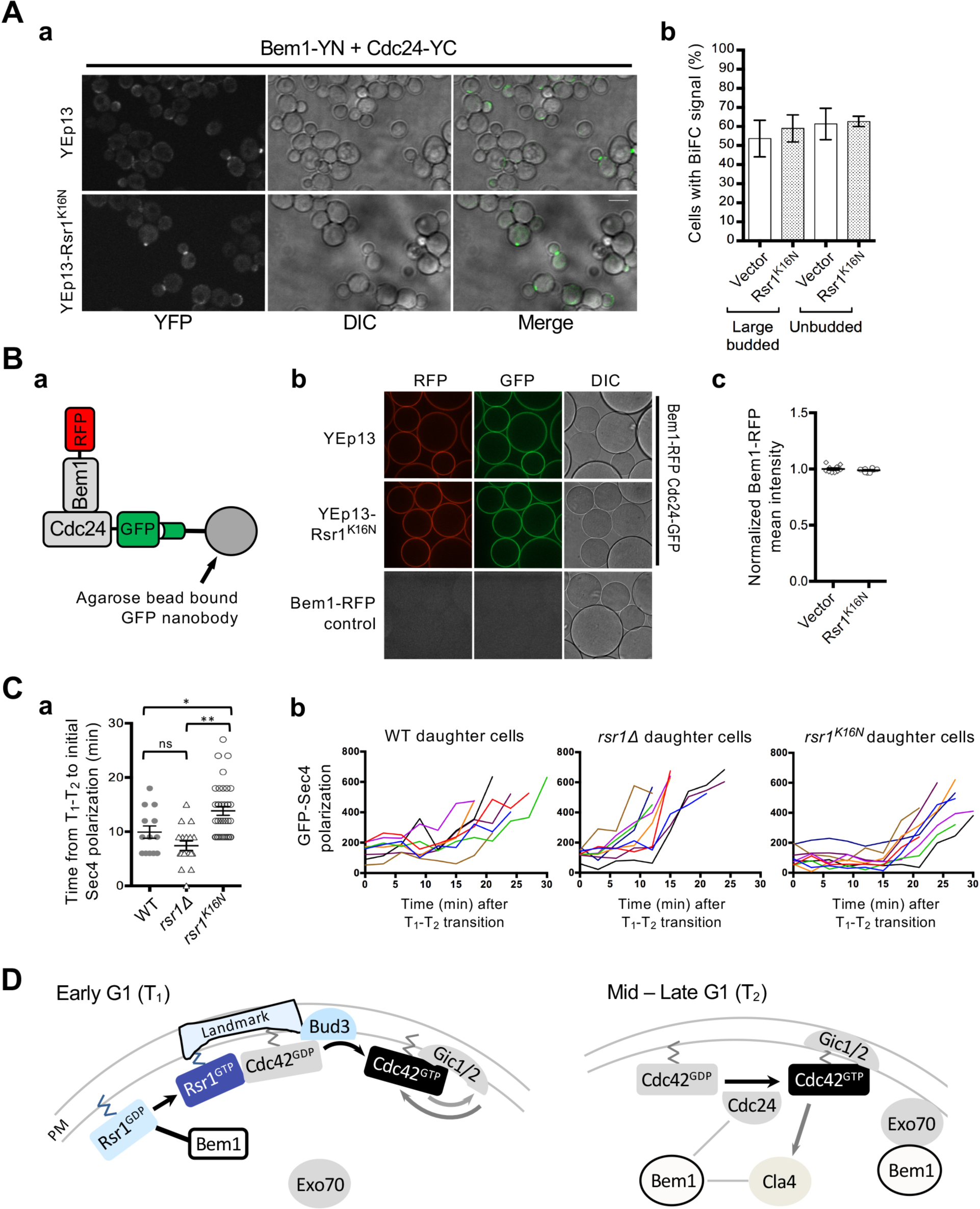
Rsr1-GDP delays Sec4 polarization but does not affect Bem1-Cdc24 interaction. (A) (a) BiFC assays in haploid *BEM1-YN CDC24-YC* carrying YEp13 or YEp13-Rsr1^K16N^. Bar, 5µm. (b) BiFC signal was quantified from three separate experiments (n=60-170 for each sample per experiment). (B) (a) Diagram of visible IP assay to test Cdc24-Bem1 interaction. (b) Images of beads from visible IP assays with marked strains and plasmids. (c) Quantification of Bem1-RFP mean intensity from individual captures of multiple beads. (C) (a) Quantification of the time interval (min) from the T_1_-T_2_ transition to initial GFP-Sec4 polarization in individual daughter cells of haploid *WT* (n=13), *rsr1*Δ (n=17), and *rsr1^K16N^* (n=36). (b) Representative graphs of GFP-Sec4 polarization in daughter cells. (D) A scheme representing a dual role of Rsr1 in spatial cue-directed polarity establishment (see text).

Next, we used ‘visible’ immunoprecipitation (VIP) assays (Katoh *et al*., 2015), which combines immunoprecipitation and microscopy, to examine whether overexpression of GDP-locked Rsr1 affects the Cdc24-Bem1 interaction. When lysates prepared from cells expressing both Cdc24-GFP and Bem1-RFP were subjected to pull-down assays using agarose beads bound to a GFP nanobody (see Materials and Methods), both Cdc24-GFP and Bem1-RFP were visible on the beads, indicating that Bem1-RFP was efficiently brought down with Cdc24-GFP. When the same strain carrying a multi-copy Rsr1^K16N^ plasmid or an empty vector were subjected to VIP assays, Bem1-RFP was recovered similarly, unlike in a control in which cell lysates containing only Bem1-RFP was used (**Figure 5B**). Collectively, these results suggest that Cdc24-Bem1 interaction is not disrupted by overexpression of Rsr1-GDP, likely because the Bem1 domains that interact with Cdc24 or Rsr1-GDP do not overlap (see **Figures**). Similarly, Bem1’s association with Rsr1-GDP may not interfere with its interaction with Cdc42. Nonetheless, Cdc42 polarization during T_2_ was delayed only in *rsr1*^*K16N*^ cells but not in *rsr1*Δ cells (see **Figure 2 & S4**), suggesting that association of Rsr1-GDP with Bem1 might hinder Cdc42 polarization, possibly by interfering with the ability of Bem1 to stimulate Cdc24 activity (Smith *et al*., 2013; Rapali *et al*., 2017).

### The interaction between Rsr1-GDP and Bem1 may ensure proper timing of polarized secretion and thus bud emergence

How does expression of the GDP-locked Rsr1 delay bud emergence? A recent report suggests that the timing of bud emergence is governed by the onset of polarized secretion (Lai *et al*., 2018). As discussed above, we found that overexpression of *rsr1^K16N^* interferes with actin-independent Exo70 localization during T_2_ and the interaction between Bem1 and Exo70. Exo70 mediates targeting and tethering of vesicles to the polarity site and is thus needed for directing polarized secretion to the incipient bud site (Boyd *et al*., 2004; He *et al*., 2007). The constitutively GDP-bound Rsr1 may continue to hold Bem1 into T_2_, resulting in delayed polarized secretion toward the bud site and consequently delayed bud emergence. To test this idea further, we compared timing of polarized secretion using the Rab GTPase GFP-Sec4, together with Whi5-RFP, in WT and *rsr1* mutants by time-lapse imaging (**Figure S6**). The onset of Sec4 polarization occurred about 10 min after the T_1_-T_2_ transition in WT cells but was delayed in *rsr1^K16N^* cells by 4 mins (**Figure 5C**), suggesting that the interaction between Bem1 and Rsr1-GDP indeed affects the timing of polarized secretion.

In summary, we show that Rsr1-GDP interacts with Bem1 in early G1 and hinders Bem1-dependent Exo70 polarization. We suggest that the Rsr1-Bem1 association may ensure the proper timing of polarized secretion for bud emergence. Based on previous reports and this new study, we propose a model whereby Rsr1 plays a dual role in polarity establishment (**Figure 5D**): in early G1, the Rsr1 GTPase cycle may be involved in linking the spatial landmark to Cdc42 polarization in haploid cells (Kang *et al*., 2014; Lee *et al*., 2015; Kang *et al*., 2018). Rsr1-GDP associates with Bem1 during T_1_ (this study), so that Bem1-mediated positive feedback may not operate until Start in haploid cells. More Rsr1 may converted to the GTP-bound form, and/or Bem1 may be modified after Start (Witte *et al.,* 2017), so that Bem1 may no longer associate with Rsr1 and can promote Cdc42 polarization and polarized secretion to the incipient bud site. While Rsr1-GDP associates with Bem1 in diploid cells as well (this study), Rsr1 appears less critical in selection of a proper bud site in diploid daughter cells: arrival of Bem1 and Cdc24 to the incipient bud site in diploid daughter cells does not depend on Rsr1 (this study) but more likely depends on the actin cytoskeleton (Yang *et al*., 1997; Harkins *et al*., 2001; Schenkman *et al*., 2002). Rsr1-GTP interacts with Cdc24 and Cdc42 (Zhang *et al*., 1995; Park *et al*., 1997; Park *et al*., 2002; Kozminski *et al*., 2003) and guides Cdc24 to the incipient bud site (Park *et al*., 2002) and may also activate Cdc24 (Shimada *et al*., 2004). Our findings highlight that the Rsr1 GTPase may have two active states that play a delicate role in coordination of spatial and temporal events leading bud emergence.

## Materials and Methods

### Strains, plasmids, and general methods

Standard methods of yeast genetics, DNA manipulation, and growth conditions were used (Guthrie and Fink, 1991). Yeast strains were grown in the appropriate synthetic medium containing 2% dextrose as a carbon source. To maintain plasmids, strains were cultured in synthetic medium lacking the appropriate nutrient(s) (e.g., SC-Ura). Yeast strains and plasmids used in this study are listed in Supplemental Tables S1 and S2, respectively, with a brief description of construction methods.

### Microscopy and image analysis

Cells were grown in synthetic medium overnight and then freshly subcultured for 3–4 hours in the same medium. Time-lapse imaging was performed essentially as previously described (Kang *et al*., 2014; Miller *et al*., 2017) using a spinning disk confocal microscope (Ultra-VIEW VoX CSU-X1 system; PerkinElmer) equipped with a 100x, 1.4 NA Plan Apochromat objective lens (Nikon), 440-, 488-, 515- and 561-nm solid-state lasers (Modular Laser System 2.0; PerkinElmer), and a backthinned electron-multiplying charge-coupled device (EM CCD) camera (ImagEM C9100-13; Hamamatsu Photonics) on an inverted microscope (Ti-E; Nikon). Images in Figures 4 and 5 were captured on the same inverted microscope but with EM CCD camera (ImageEM X2 C9100-23B; Hamamatsu Photonics). For most time-lapse imaging, images were captured (9 z stacks, 0.3 µm step for haploid cells; 11 z stacks, 0.4 µm step for diploid cells) every 3 or 5 min using cells mounted on an agarose slab at either room temperature or 30°C, as indicated in figure legend.

Image processing and analyses were performed using ImageJ (National Institutes of Health). Fluorescent images in figures are generated using maximum intensity projections of z stacks, whereas a single middle z-section is shown for DIC images. To quantify the Whi5-GFP or Whi5-RFP signal in the nucleus at each time point, a circular ROI that included the Whi5 signal in the nucleus was used to measure the intensity from summed intensity projection images after background subtraction. The T_1_-T_2_ transition was marked when the Whi5 intensity in the nucleus was about 50% of its peak level (Skotheim *et al*., 2008; Donic *et al*., 2011). The duration time of T_1_ was considered from the onset of cytokinesis (estimated when PBD-RFP level was the lowest (Okada *et al*., 2013), which was about 3-5 min after the nuclear entry of Whi5 entry at 30°C (Di Talia *et al*., 2007; Lee *et al*. 2015) until the T_1_-T_2_ transition. The duration time of T_2_ was determined from the T_1_-T_2_ transition until bud emergence (which was estimated from PBD level and DIC images).

To quantify the Bem1-RFP signal, average intensity projections were analyzed after background subtraction, and an ROI was drawn around the daughter cell. The Bem1-RFP integrated density values were obtained for each time point captured over the G1 phase using a threshold method, as previously described (Okada *et al*., 2013; Lee *et al*., 2015). The peak Bem1-RFP values during T_1_ or T_2_ were normalized to the average Bem1-RFP peak level in WT cells during T_1_ or T_2_, respectively (Figure 1C).

The PBD-RFP cluster in daughter cells was quantified using a threshold method (Okada *et al*., 2013; Lee *et al*., 2015) from average intensity projection images of five selected z-sections after background subtraction. The PBD-RFP integrated density values were obtained for each time point captured over the G1 phase. The peak PBD-RFP level during T_2_ was determined and the time from the T_1_-T_2_ transition until the peak PBD-RFP level was calculated for each individual daughter cell (Figure 2Bb & S4Ba). Sec4-GFP polarization was analyzed similarly using a threshold method (Okada *et al*., 2013; Lee *et al*., 2015), and the time from the T_1_-T_2_ transition until detection of initial Sec4-GFP polarization at the incipient bud site was determined for each individual daughter cell (Figure 5C)

To quantify BiFC signals, summed intensity projections were analyzed after background subtraction. A fluorescence threshold was set above background that selected fluorescent pixels at the division site of large budded and unbudded cells. The mean grey value of the YFP signal above the threshold at the division site of each cell was measured. Values were normalized to the average BiFC signal intensity with Bem1-YN in large-budded or unbudded cells, respectively (Figure 3B). To quantify the percent of cells with positive BiFC signals (Figure 4B and 5A), summed intensity projections were analyzed after background subtraction. A fluorescence threshold was set above background that selected fluorescent pixels at the division site of large budded and unbudded cells. The percent of cells with a BiFC signal above the threshold was determined from three independent experiments (Figure 4Bb and 5Ab).

Cells with polarized Exo70-RFP were quantified at different cell cycle stages based on localization of Whi5-GFP and DIC images: large budded cells with Whi5-GFP in the nucleus (late M); unbudded cells with Whi5-GFP in the nucleus (T_1_); unbudded cells without Whi5-GFP (T_2_); and small budded cells. Summed intensity projections of Exo70-RFP z-stack images were created after background subtraction. A fluorescence threshold was set above background that selected fluorescent pixels at the division site in late M and T_1_ cells, or at the incipient bud site in T_2_ cells, or at the tips of the growing buds in small budded cells. The percent of cells with a signal above the threshold was determined from three independent experiments (Figure 4Ab).

To quantify bead bound Bem1-RFP from visible immunoprecipitation assays, summed intensity projections of z-stacks were analyzed after background subtraction. Images of the Bem1-RFP alone control was used to determine a threshold that selected fluorescent pixels above background. The same threshold was applied to all images and the mean grey value of all pixels above the set threshold was measured (Figure 5Bc).

### FRAP analysis

To perform FRAP experiments, images were captured at a single z section on a gelatin slab at 22°C using the photokinesis unit on the Ultra-VIEW VoX confocal system (see above), as previously described (Miller et al., 2017; Kang et al., 2018). Prior to beginning each FRAP experiment, a z-stack image was taken with the 561-nm laser to examine the Whi5-mCherry signal and select cells in T_1_ or T_2_. The middle focal plane of cells was chosen to bleach. After collecting 5 pre-bleach images, selected ROI’s were bleached to <50% of the original fluorescence intensity. Post-bleach images were acquired for a duration long enough so that the recovery curve reached a plateau. After background subtraction and correcting for photobleaching, the data were normalized to the mean pre-bleach intensity of the ROI set to 100% and the intensity just after bleaching set to 0% so that FRAP curves show the percentage of recovery. To reduce noise, the intensity of every 3 consecutive post-bleach time points was averaged. The intensity data were plotted and fitted using the exponential equation y = m_1_+m_2_ * exp(-m_3_ * X), where m_3_ is the off-rate, using Prism 6 (GraphPad Software). The half-time of recovery was calculated using the equation t_1/2_ = (ln2)/m_3_.

### Bimolecular fluorescence complementation (BiFC) assays

YC fusions of WT or mutant Rsr1 proteins and VC-Cdc42 were expressed from their chromosomal loci, as previously described (Kang *et al*., 2010). YN fusions of WT or mutant Bem1 proteins were expressed either using multicopy plasmids or from the chromosome. Cdc24-VC or Exo70-VC were expressed from each chromosomal locus (see Tables S1 and S2). Each combination of YC (or VC) and YN fusion proteins were expressed in haploid cells (unless indicated otherwise) and subjected to microscopy (see below). Since the same split site (154/155) was used to generate both YFP and Venus truncated forms for BiFC, YC or VC fusions were tested in combination with a YN fusion.

For BiFC assays, cells were grown in the appropriate synthetic medium overnight and then freshly subcultured for 3-4 hours in the same medium prior to imaging. Cells were mounted on an agarose slab containing the same medium, and static images were captured (5 z-stacks, 0.3 µm step for haploid cells; 5 z-stacks, 0.4 µm step for diploid cells) using a spinning disk confocal microscope (see above) at room temperature. Time-lapse images of haploid cells expressing YC-Rsr1^K16N^ carrying pRS426-Bem1-YN were captured similarly except every 10 min (Figure 1Ab).

### LatA treatment

Cells were grown in SC-LEU medium (to maintain YEp13 plasmids) overnight and then freshly subcultured for 3–4 hours in the same medium at 30°C. Cells were harvested and treated with 100 µM LatA for 10 min or mock-treated with DMSO before imaging using a spinning disk confocal microscope (see above).

### Visible Immunoprecipitation (VIP) assay

DLY13038 (*CDC24-GFP BEM1-RFP*) carrying YEp13-RSR1^K16N^ or YEp13 were grown in SC-LEU, and HPY3336 (*BEM1-RFP*) was grown in YPD medium overnight. These cells were then freshly subcultured for 3–4 hours in the same medium at 30°C until mid-log phase. A total of 44 OD_600_ units of cells were harvested and cell lysates were prepared using buffer VII (200mM KCl, 1% Triton X-100, 10% glycerol, 1mM MgCl_2_, 50mM HEPES pH 7.6, 1mM EGTA) along with a cocktail of protease inhibitors. Crude cell lysates were centrifuged for 12 min at 10,000 g, and the supernatant (S10 fraction) was used for subsequent assays. This S10 fraction was then diluted with an equal volume of buffer VII lacking KCl and Triton X-100 and incubated with 10µL of GFP-Trap beads (gta-10, Chromotek) for 1 hour at 4°C by rocking. The beads were then washed 4 times using a wash buffer (50mM KCl, 0.1% Triton X-100, 10% glycerol, 1mM MgCl_2_, 50mM HEPES pH 7.6, 1mM EGTA, 1 mM dithiothreitol). Beads were resuspended in a small volume of the same buffer and immediately mounted on an agarose slab, and static images were captured (5 z-stacks, 0.5µm step) using a spinning disk confocal microscope (see above) with a 40x, NA 1.3 Plan Fluor oil objective lens (Nikon) at room temperature.

### Statistical analysis and graph presentation

Data analysis was performed using Prism 6 (GraphPad Software). Graphs in figures show Mean (horizontal lines) ± SEM (error bars) unless indicated otherwise. The bar graphs of FRAP data shows median as a line, quartiles, maximum, and minimum (Figure 1Ba). A two-tailed student’s t test was performed to determine statistical differences between two sets of data: ns (not significant) for p ≥ 0.05; *p < 0.05; **p < 0.01; and ***p < 0.001. Pearson correlation analysis was used to determine the strength of a linear association between T_2_ length and the peak Cdc42-GTP arrival time after T_1_-T_2_ transition.

## Acknowledgements

We are grateful to E. Bi, D. Lew, W.-K. Huh, and R. Ruggieri for providing yeast strains and plasmids; and P. J. Kang for discussion and comments on the manuscript. This work was supported by a research grant (R01-GM114582) from the NIH/NIGMS to HOP. KEM was partly supported by the Seilhamer Fellowship from The Jeffrey J. Seilhamer Cancer Foundation.

## Abbreviations

GTPase: guanosine triphosphatases
GEF: guanine nucleotide exchange factor
GAP: GTPase-activating protein
PAK: p21-activated kinase
PBD: p21-binding domain
WT: wild-type
FRAP: fluorescence recovery after photobleaching
BiFC: bimolecular fluorescence complementation
YFP: yellow fluorescent protein
YN: N-terminal half of YFP
YC: C-terminal half of YFP
LatA: Latrunculin A.

