## Supplemental Figures and Tables for "Temporal regulation of cell polarity via the interaction of the Ras GTPase Rsr1 and the scaffold protein Bem1"

**Figure Legends**

**Figure S1**. Bem1 localization in haploid cells during G1. Time-lapse images of Bem1-RFP and Whi5-GFP in (a) WT and (b) *rsr1*$^{K16N}$ cells at 30°C; Bem1-GFP and PBD-RFP in WT cells at 22°C (c); Bem1-RFP and Cdc24-GFP in (d) WT and (e) *rsr1*Δ cells at 22°C. Numbers indicate time (min) relative to the onset of cytokinesis (t=0). Asterisks (a & b) mark $T_1$-$T_2$ transition in daughter cells. An arrow in (d) points to weak Cdc24-GFP signal at the division site. Bars, 3 μm.

**Figure S2**. Bem1 localization in diploid cells during G1. Time-lapse images of Bem1-RFP and Whi5-GFP in (a) *WT*, (b) *rsr1*Δ, and (c) *rsr1*$^{K16N}$ cells at 22°C. Numbers indicate time (min) relative to the onset of cytokinesis (t=0). Asterisks mark $T_1$-$T_2$ transition in daughter cells. Yellow and red arrows mark localized Bem1-RFP signal during $T_1$ and at the incipient bud site during $T_2$, respectively. Bars, 3 μm.

**Figure S3.** Bem1 and Cdc24 localization in diploid cells during G1. (A) Time-lapse images of (a) Cdc24-GFP and Whi5-RFP in WT cells; and Bem1-RFP and Cdc24-GFP in (b) WT and (c) *rsr1*Δ cells at 22°C. Numbers indicate time (min) relative to the onset of cytokinesis (t=0). An asterisk marks $T_1$-$T_2$ transition in the daughter cell (a). Yellow and red arrows mark localized Cdc24 or Bem1 signals during $T_1$ and at the incipient bud site during $T_2$, respectively (a-c). Bars, 3 μm. (B) Localization pattern (%) of Bem1 (b) or Cdc24 (c), marked in red in (a), during $T_1$ and $T_2$ (Whi5 in yellow) in daughter cells is summarized from time-lapse images of WT (n=45), *rsr1*$^{K16N}$ (n=10), and *rsr1*Δ cells (n=20).

**Figure S4.** The second phase of Cdc42 polarization is delayed in diploid cells expressing GDP-locked Rsr1. (A) Time-lapse images of PBD-RFP and Whi5-GFP in (a) WT and (b) *rsr1*$^{K16N}$ and (c) *rsr1*Δ cells at 22°C. Numbers indicate time (min) relative to the onset of cytokinesis (t=0). Asterisks mark $T_1$-$T_2$ transition in daughter cells. Bars, 3 μm. (B) (a) Quantification of the time (min) from $T_1$-$T_2$ transition until Cdc42-GTP reached a maximum level during $T_2$ in WT (n=11), *rsr1*Δ (n=27), and *rsr1*$^{K16N}$ (n=11) daughter cells. Values are shown for each individual cell. (b) Representative graphs showing Cdc42-GTP polarization over time (min) from the onset of cytokinesis (t=0) in daughter cells. Each line shows Cdc42 polarization in a single daughter cell marked with the $T_1$-$T_2$ transition point (arrow) and when Cdc42-GTP level peaked during $T_2$ (arrowhead). (C) (a) Length of $T_1$ and $T_2$ (min) in each daughter cell (WT, n =11; *rsr1*Δ, n =27; and *rsr1*$^{K16N}$, n =11). (b) Correlation analysis of $T_2$ length and peak Cdc42-GTP arrival time after the $T_1$-$T_2$ transition in *rsr1*$^{K16N}$ cells (n=11).

**Figure S5.** BiFC assays of additional *bem1* mutants in haploids and WT Bem1 in diploids. (A) BiFC assays in haploid cells expressing YC-Cdc42, YC-Rsr1$^{G12V}$, or YC-Rsr1$^{K16N}$ and carrying (a) pRS426-Bem1-YN, (b) pRS426- bem1$^{\Delta 1\text{-}147}$-YN, (c) pRS426- bem1$^{\Delta 148\text{-}280}$-YN, (d) pRS426-bem1$^{K482A}$-YN, or (e) pRS426-bem1$^{\Delta 412\text{-}551}$-YN. (B) Diagram of WT and mutant Bem1 proteins. Regions deleted are marked with delta (Δ). An asterisk (*) marks the K482A mutation. (C) BiFC assays in diploid cells expressing Bem1-YN and YC-Rsr1$^{K16N}$ or YC-Rsr1$^{G12V}$. Bars, 5μm.

**Figure S6.** Sec4 polarization in haploid cells. Time-lapse images of GFP-Sec4 and Whi5-RFP in (a) WT, (b) *rsr1*$^{K16N}$*,* and (c) *rsr1*Δ cells at 30°C. Numbers indicate time (min) relative to $T_1$-$T_2$ transition (marked with an asterisk; t=0) in daughter cells. Green arrowheads point the first appearance of polarized GFP-Sec4. Bars, 3 μm.

### Figure S1

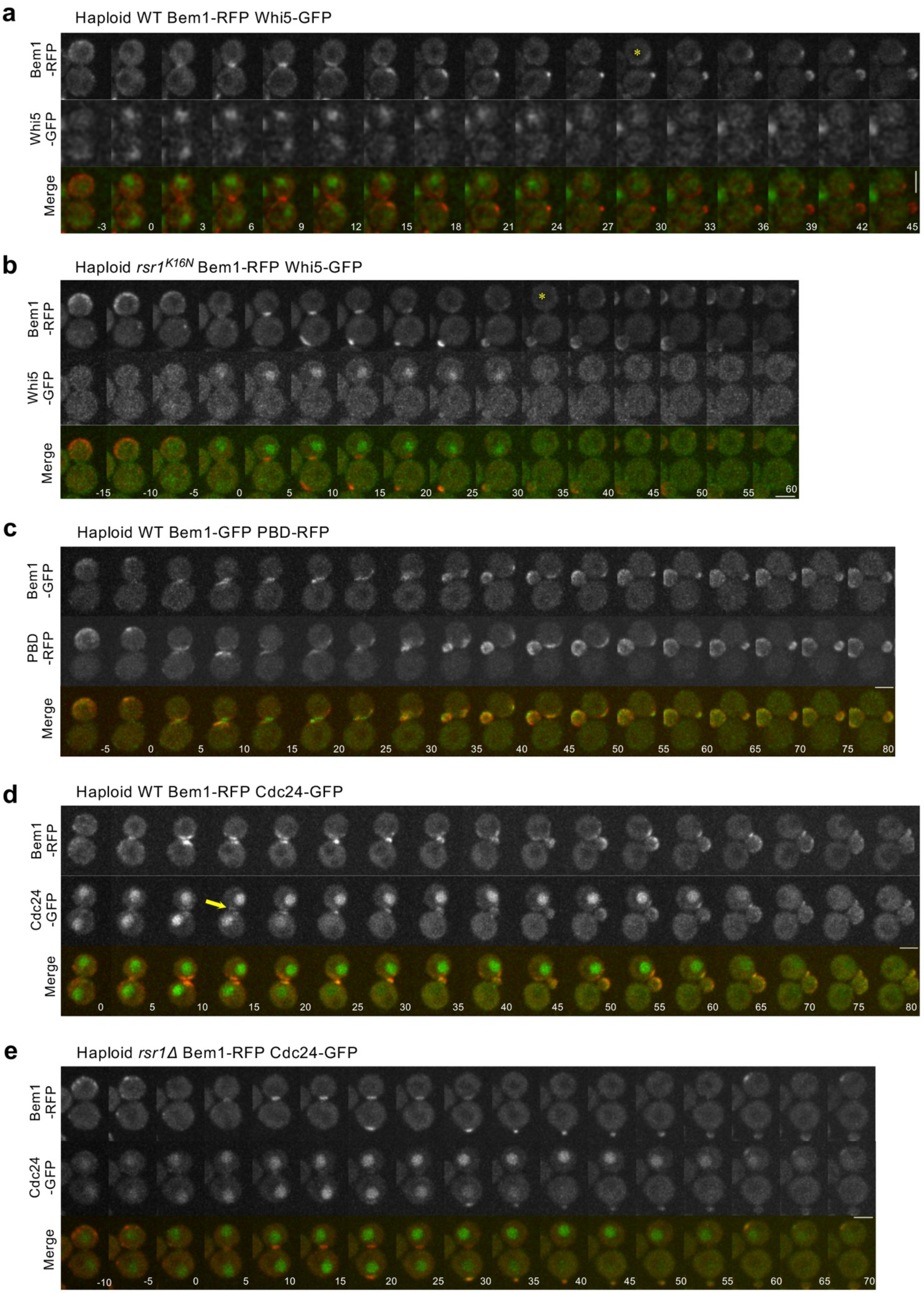
a
Haploid WT Bem1-RFP Whi5-GFP
Bem1 -RFP
Whi5 -GFP
Merge
-3 0 3 6 9 12 15 18 21 24 27 30 33 36 39 42 45
b
Haploid rsr1K16N Bem1-RFP Whi5-GFP
Bem1 -RFP
Whi5 -GFP
Merge
-15 -10 -5 0 5 10 15 20 25 30 35 40 45 50 55 60
c
Haploid WT Bem1-GFP PBD-RFP
Bem1 -GFP
PBD -RFP
Merge
-5 0 5 10 15 20 25 30 35 40 45 50 55 60 65 70 75 80
d
Haploid WT Bem1-RFP Cdc24-GFP
Bem1 -RFP
Cdc24 -GFP
Merge
0 5 10 15 20 25 30 35 40 45 50 55 60 65 70 75 80
e
Haploid rsr1Δ Bem1-RFP Cdc24-GFP
Bem1 -RFP
Cdc24 -GFP
Merge
-10 -5 0 5 10 15 20 25 30 35 40 45 50 55 60 65 70

### Figure S2

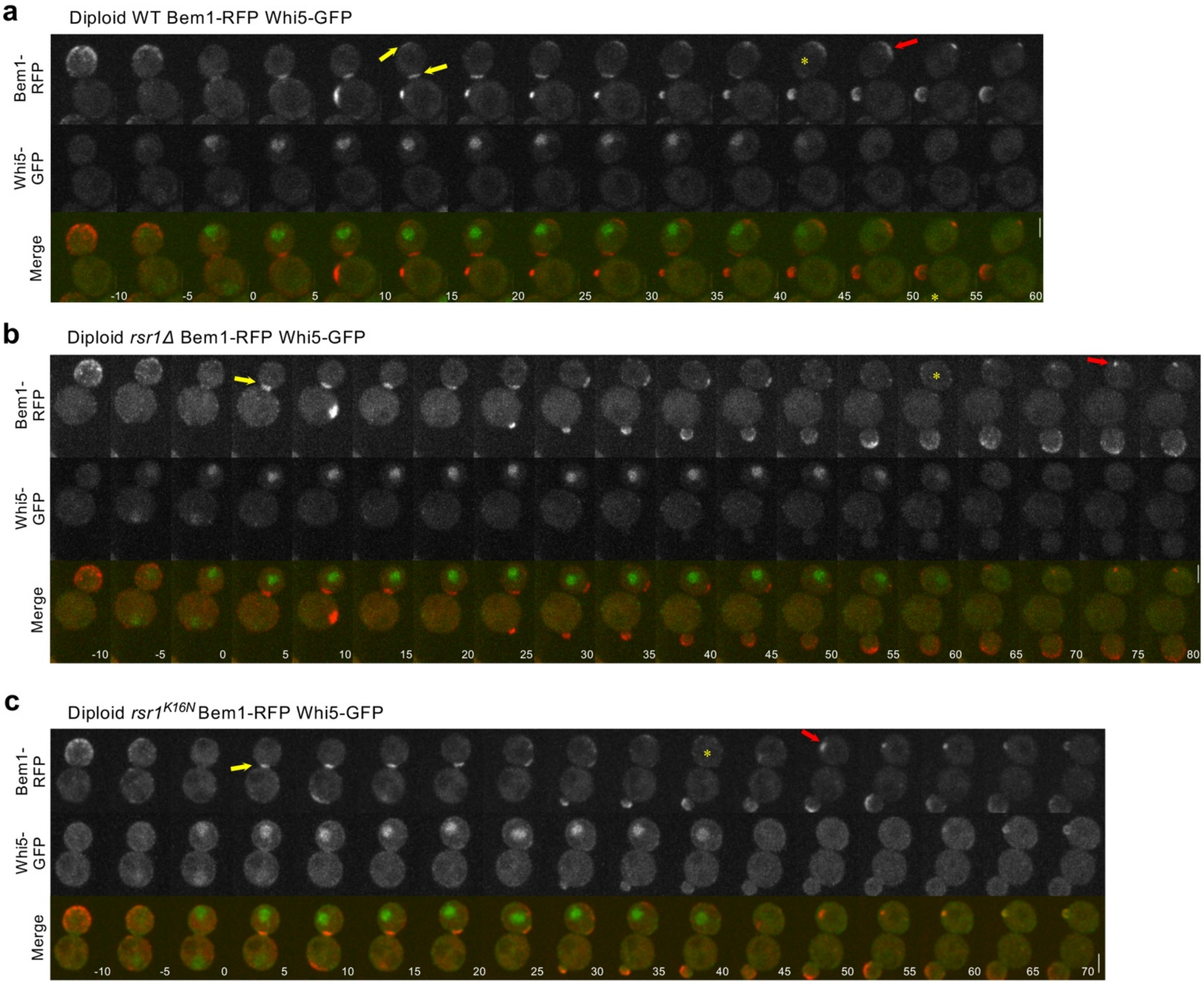
a
Diploid WT Bem1-RFP Whi5-GFP
Bem1-RFP
Whi5-GFP
Merge
-10
-5
0
5
10
15
20
25
30
35
40
45
50
55
60
b
Diploid rsr1Δ Bem1-RFP Whi5-GFP
Bem1-RFP
Whi5-GFP
Merge
-10
-5
0
5
10
15
20
25
30
35
40
45
50
55
60
65
70
75
80
c
Diploid rsr1K16N Bem1-RFP Whi5-GFP
Bem1-RFP
Whi5-GFP
Merge
-10
-5
0
5
10
15
20
25
30
35
40
45
50
55
60
65
70

### Figure S3

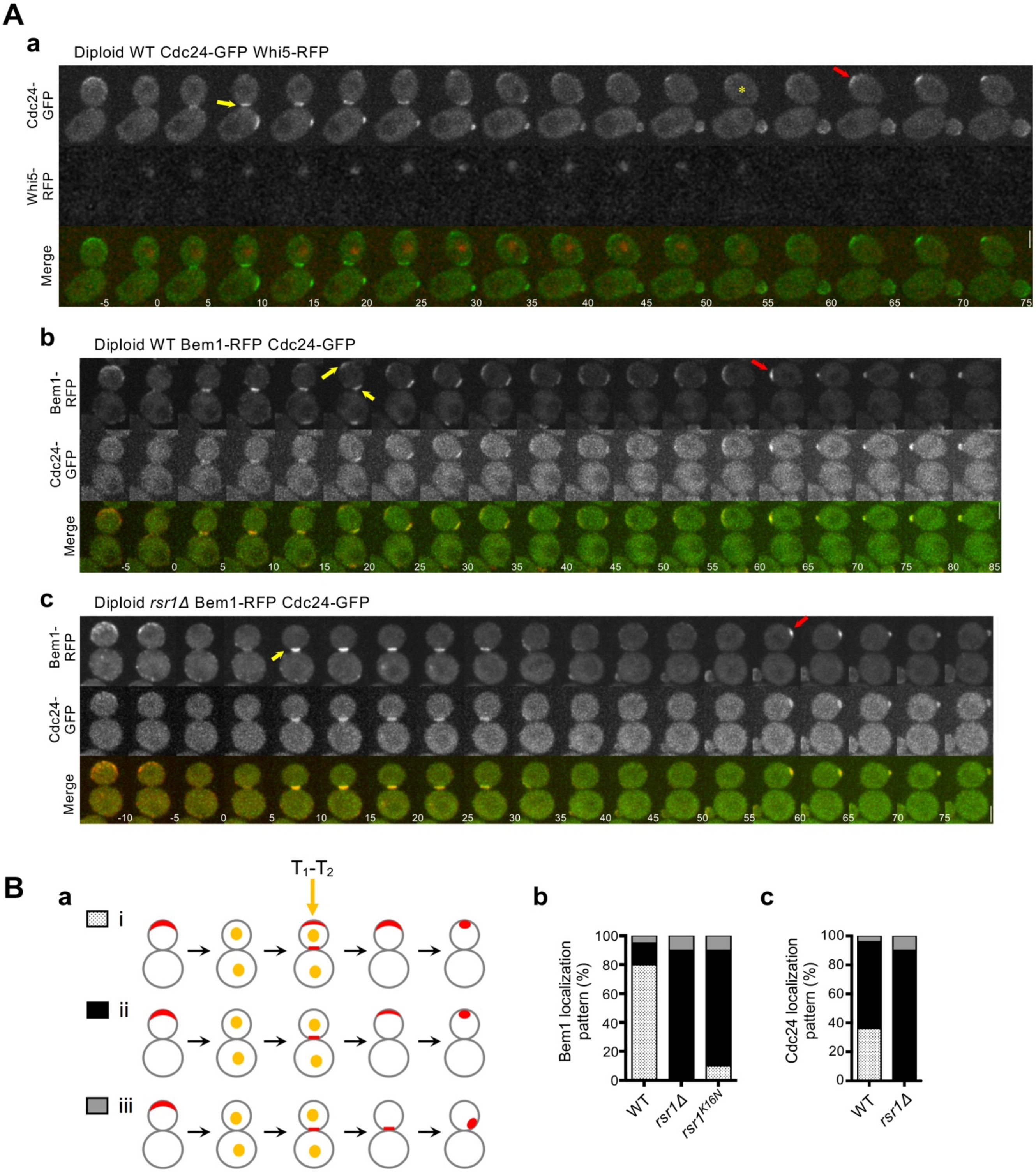
A
a
Diploid WT Cdc24-GFP Whi5-RFP
Cdc24-GFP
Whi5-RFP
Merge
-5 0 5 10 15 20 25 30 35 40 45 50 55 60 65 70 75
b
Diploid WT Bem1-RFP Cdc24-GFP
Bem1-RFP
Cdc24-GFP
Merge
-5 0 5 10 15 20 25 30 35 40 45 50 55 60 65 70 75 80 85
c
Diploid rsr1Δ Bem1-RFP Cdc24-GFP
Bem1-RFP
Cdc24-GFP
Merge
-10 -5 0 5 10 15 20 25 30 35 40 45 50 55 60 65 70 75
B
a
T1-T2
i
ii
iii
b
Bem1 localization pattern (%)
0 20 40 60 80 100
WT
rsr1Δ
rsr1K16N
c
Cdc24 localization pattern (%)
0 20 40 60 80 100
WT
rsr1Δ

### Figure S4

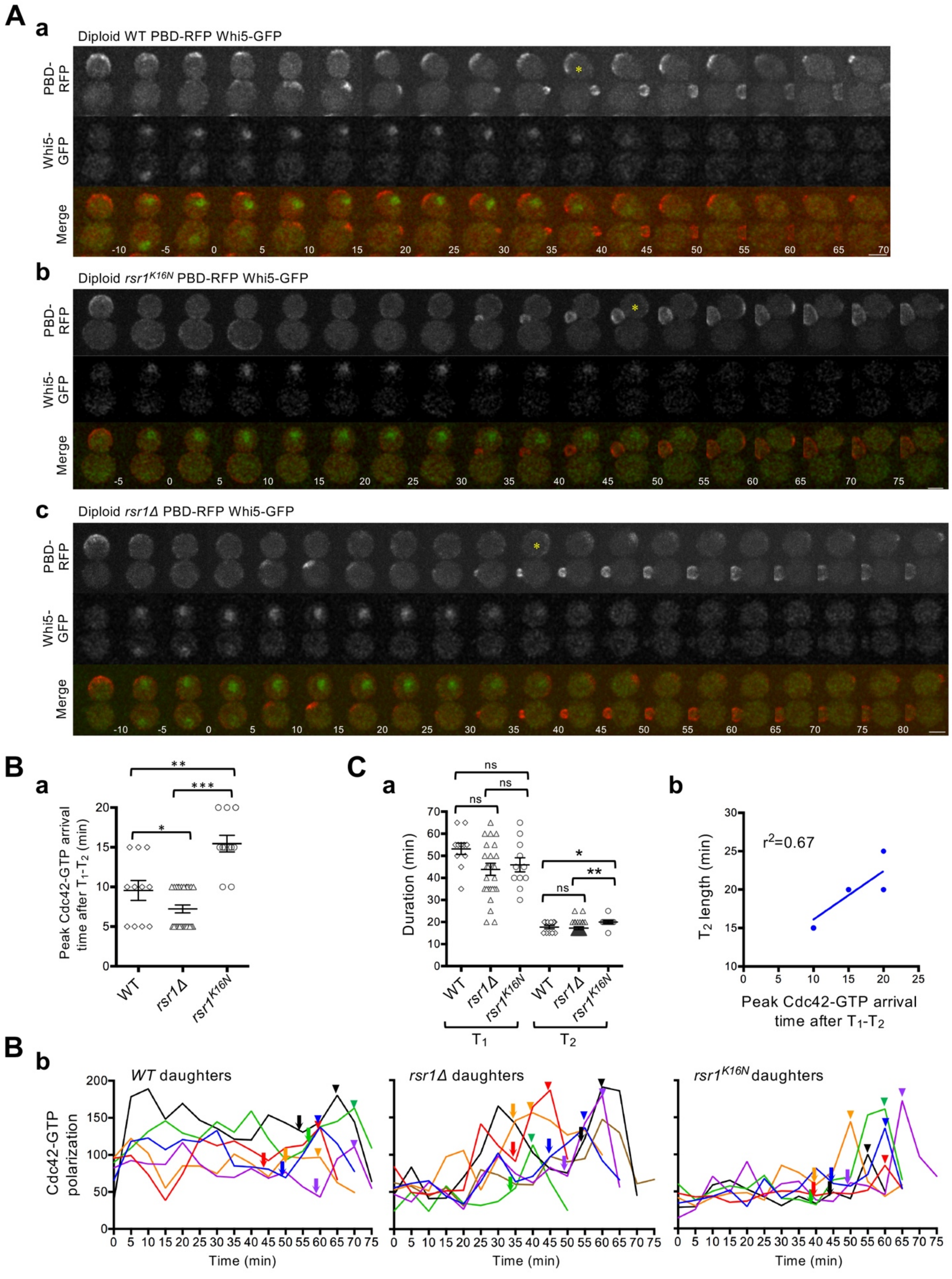

A
a
Diploid WT PBD-RFP Whi5-GFP
PBD-RFP
Whi5-GFP
Merge
-10 -5 0 5 10 15 20 25 30 35 40 45 50 55 60 65 70
b
Diploid rsr1K16N PBD-RFP Whi5-GFP
PBD-RFP
Whi5-GFP
Merge
-5 0 5 10 15 20 25 30 35 40 45 50 55 60 65 70 75
c
Diploid rsr1Δ PBD-RFP Whi5-GFP
PBD-RFP
Whi5-GFP
Merge
-10 -5 0 5 10 15 20 25 30 35 40 45 50 55 60 65 70 75 80
B
a
Peak Cdc42-GTP arrival time after T1-T2 (min)
WT
rsr1Δ
rsr1K16N
C
a
Duration (min)
WT
rsr1Δ
rsr1K16N
T1
T2
ns
b
r2=0.67
T2 length (min)
Peak Cdc42-GTP arrival time after T1-T2
B
b
WT daughters
rsr1Δ daughters
rsr1K16N daughters
Cdc42-GTP polarization
Time (min)

### Figure S5

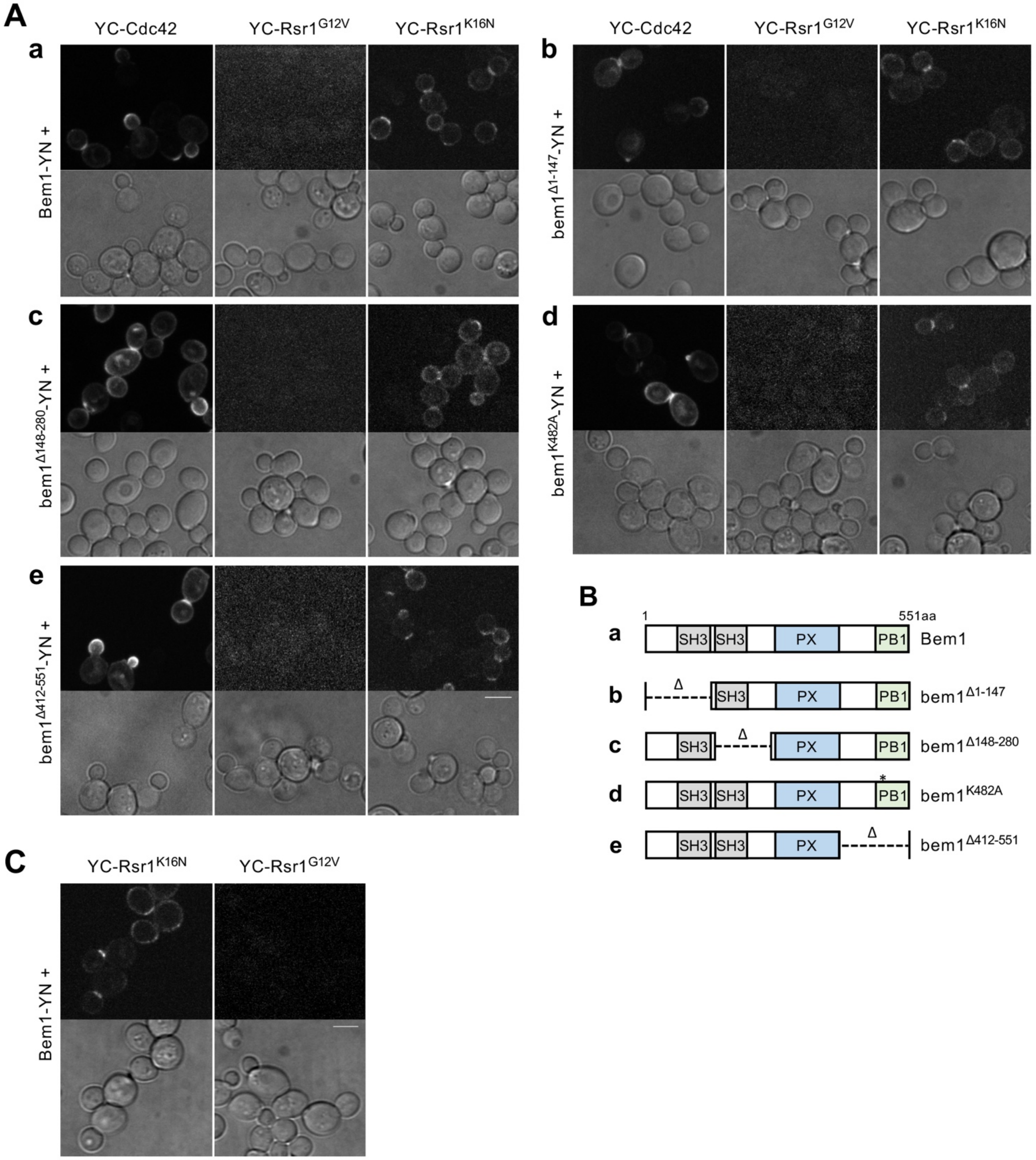

A
YC-Cdc42
YC-Rsr1G12V
YC-Rsr1K16N
a
Bem1-YN +
b
bem1Δ1-147-YN +
c
bem1Δ148-280-YN +
d
bem1K482A-YN +
e
bem1Δ412-551-YN +
B
1
551aa
SH3
PX
PB1
Bem1
Δ
bem1Δ1-147
bem1Δ148-280
*
bem1K482A
bem1Δ412-551
C
YC-Rsr1K16N
YC-Rsr1G12V
Bem1-YN +

#### Figure S6

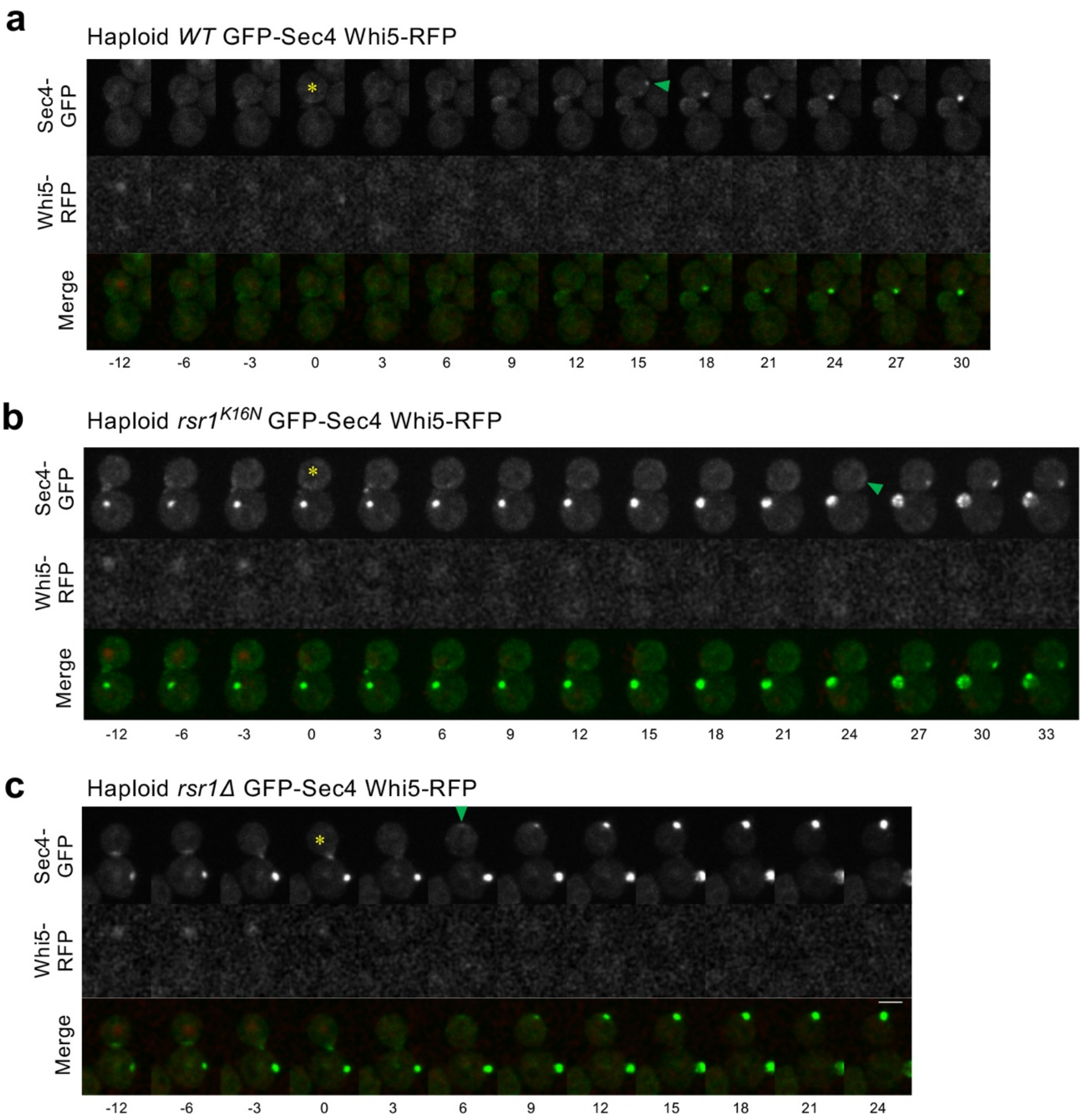
a
Haploid WT GFP-Sec4 Whi5-RFP
Sec4-GFP
Whi5-RFP
Merge
-12
-6
-3
0
3
6
9
12
15
18
21
24
27
30
b
Haploid rsr1K16N GFP-Sec4 Whi5-RFP
Sec4-GFP
Whi5-RFP
Merge
-12
-6
-3
0
3
6
9
12
15
18
21
24
27
30
33
c
Haploid rsr1Δ GFP-Sec4 Whi5-RFP
Sec4-GFP
Whi5-RFP
Merge
-12
-6
-3
0
3
6
9
12
15
18
21
24

**Table S1. Yeast strains used in this study**

| Strain | Relevant Genotype[a] | Source/Comments |
|---|---|---|
| YEF473A | **a** *his3-Δ200 leu2-Δ1 lys2-801 trp1-Δ63 ura3-52* | Bi and Pringle, 1996 |
| HPY1197 | **a** *VC-CDC42-KAN* | Kang *et al.,* 2010 |
| HPY1213 | **a** *YFP*$^{C}$*-RSR1-TRP1* | Kang *et al.,* 2010 |
| HPY1522 | **a** *YFP*$^{C}$*-rsr1*$^{K16N}$*-TRP1* | Kang *et al.,* 2010 |
| HPY1552 | **a** *YFP*$^{C}$*-rsr1*$^{G12V}$*-TRP1* | Kang *et al.,* 2010 |
| HPY3340 | **a** *VC-CDC42-KAN BEM1-YFP*$^{N}$*-URA3* | This Study[b] |
| HPY3441 | **a** *YFP*$^{C}$*-RSR1-TRP1 BEM1-YFP*$^{N}$*-URA3* | This Study[b] |
| HPY3442 | **a** *YFP*$^{C}$*-rsr1*$^{K16N}$*-TRP1 BEM1-YFP*$^{N}$*-URA3* | This Study[b] |
| HPY3443 | **a** *YFP*$^{C}$*-rsr1*$^{G12V}$*-TRP1 BEM1-YFP*$^{N}$*-URA3* | This Study[b] |
| HPY3444 | **a** *VC-CDC42-KAN bem1*$^{\Delta 281\text{-}345}$*-YFP*$^{N}$*-URA3* | This Study[b] |
| HPY3445 | **a** *YFP*$^{C}$*-RSR1-TRP1 bem1*$^{\Delta 281\text{-}345}$*-YFP*$^{N}$*-URA3* | This Study[b] |
| HPY3446 | **a** *YFP*$^{C}$*-rsr1*$^{K16N}$*-TRP1 bem1*$^{\Delta 281\text{-}345}$*-YFP*$^{N}$*-URA3* | This Study[b] |
| HPY3447 | **a** *YFP*$^{C}$*-rsr1*$^{G12V}$*-TRP1 bem1*$^{\Delta 281\text{-}345}$*-YFP*$^{N}$*-URA3* | This Study[b] |
| HPY3448 | **a** *VC-CDC42-KAN bem1*$^{\Delta 345\text{-}408}$*-YFP*$^{N}$*-URA3* | This Study[b] |
| HPY3449 | **a** *YFP*$^{C}$*-RSR1-TRP1 bem1*$^{\Delta 345\text{-}408}$*-YFP*$^{N}$*-URA3* | This Study[b] |
| HPY3450 | **a** *YFP*$^{C}$*-rsr1*$^{K16N}$*-TRP1 bem1*$^{\Delta 345\text{-}408}$*-YFP*$^{N}$*-URA3* | This Study[b] |
| HPY3451 | **a** *YFP*$^{C}$*-rsr1*$^{G12V}$*-TRP1 bem1*$^{\Delta 345\text{-}408}$*-YFP*$^{N}$*-URA3* | This Study[b] |
| HPY2671 | α *WHI5-GFP-TRP1 PBD-tdTomato-URA3* | Lee *et al.,* 2015 |
| HPY2669 | α *rsr1Δ::URA3 WHI5-GFP-KAN PBD-tdTomato-URA3* | Lee *et al.,* 2015 |
| HPY3296 | **a** *BEM1-GFP-LEU2 WHI5-mCherry-hph* | This Study[c] |
| HPY3300 | **a** *rsr1Δ::TRP1 BEM1-GFP-LEU2 WHI5-mCherry-hph* | This Study[c] |
| HPY3218 | α *rsr1Δ::URA3 rsr1*$^{K16N}$*-TRP1 WHI5-GFP-KAN PBD-tdTomato-URA3* | This Study[d, e] |
| HPY3190 | **a**/α *WHI5-GFP-KAN PBD-tdTomato-URA3* | This Study[d] |
| HPY3259 | **a**/α *rsr1Δ::URA3 rsr1*$^{K16N}$*-TRP1 WHI5-GFP-KAN PBD-tdTomato-URA3* | This Study[d, e] |

| | | |
|---|---|---|
| HPY3331 | **a**/α *rsr1Δ::URA3 WHI5-GFP-KAN PBD-tdTomato-URA3* | This Study[d] |
| DLY9875 | α *PBD-tdTomato-URA3 BEM1-GFP-LEU2* | Daniel Lew |
| DLY13038 | **a** *CDC24-GFP-TRP1 BEM1-tdTomato-HIS3* | Daniel Lew |
| HPY3349 | **a**/α *CDC24-GFP-TRP1 BEM1-tdTomato-HIS3* | This Study |
| HPY3367 | **a**/α *rsr1Δ::URA3 BEM1-tdTomato-HIS3 CDC24-GFP-LEU2* | This Study |
| HPY3370 | **a**/α *rsr1Δ::URA3 rsr1*$^{K16N}$*-TRP1 BEM1-tdTomato-HIS3 WHI5-GFP-KAN* | This Study[d, e] |
| HPY3342 | α *rsr1Δ::URA3 rsr1*$^{K16N}$*-TRP1 BEM1-GFP-LEU2 WHI5-mCherry-hph* | This Study[c, e] |
| HPY3426 | **a** *GFP-SEC4-URA3 WHI5-mCherry-hph* | This Study[c] |
| HPY3427 | α *rsr1Δ::TRP1 GFP-SEC4-URA3 WHI5-mCherry-hph* | This Study[e] |
| HPY3430 | **a** *rsr1Δ::URA3 rsr1*$^{K16N}$*-TRP GFP-SEC4-URA3 WHI5-mCherry-hph* | This Study[c, e] |
| HPY3319 | **a** *BEM1-tdTomato-HIS3 WHI5-GFP-TRP1* | This Study[d] |
| HPY3461 | α *EXO70-tdTomato-KAN WHI5-GFP-TRP1* | This Study[d] |
| HPY3473 | **a** *BEM1-YFP*$^{N}$*-URA3 EXO70-VC-KAN* | This Study[f] |
| HPY3483 | **a** *BEM1-YFP*$^{N}$*-URA3 CDC24-VC-KAN* | This Study[f] |
| HPY3368 | **a** *rsr1Δ::URA3 rsr1*$^{K16N}$*-TRP1 WHI5-GFP-KAN BEM1-tdTomato-HIS3* | This Study [d, e] |
| HPY3347 | **a** *rsr1Δ::URA3 CDC24-GFP-TRP1 BEM1-tdTomato-HIS3* | This Study |
| HPY3380 | **a**/α *BEM1-tdTomato-HIS3 WHI5-GFP-TRP1* | This Study[d] |
| HPY3231 | **a**/α *WHI5-mCherry-hph CDC24-GFP-TRP1* | This Study[c] |
| HPY3467 | **a**/α *rsr1Δ::URA3 BEM1-tdTomato-HIS3 WHI5-GFP-TRP1* | This Study[d] |
| HPY3336 | **a** *BEM1-tdTomato-HIS3* | This Study |

[a] All strains are congenic to YEF473A. The original strains and plasmids expressing Gic2-PBD-tdTomato were previously described (Tong *et al*., 2007) (kind gifts from E. Bi, University of Pennsylvania). The original strains and plasmids expressing Bem1-GFP (Kozubowski *et al.,* 2008), Bem1-tdTomato (Howell *et al.,* 2012), and GFP-Sec4 (Chen *et al*., 2012) were previously described (gifts from D. Lew, Duke University).

[b] pRS306 plasmid carrying BEM1-YFP$^{N}$ or bem1$^{\Delta 281\text{-}345}$-YFP$^{N}$ or bem1$^{\Delta 345\text{-}408}$-YFP$^{N}$ was linearized with *Stu*I and integrated at the *ura3* locus.

[c] *WHI5-mCherry* was constructed by the C-terminal tagging method described in Longtine *et al*. (1998) using pBS35 (mCherry, hygromycin-B selection; a gift from Yeast Resource Center, University of Washington, Seattle, WA), replacing the endogenous *WHI5* (Miller *et al.,* 2017).
[d] *WHI5-GFP* was constructed by the C-terminal tagging method described in Longtine *et al*., (1998), using pFA6a-GFP-TRP1 or pFA6a-GFP-kanMX6 replacing the endogenous *WHI5* (Kang *et al*., 2014).
[e] rsr1$^{K16N}$-TRP1 contains a YFP tag at the N terminus (Park *et al.,* 2002). Since the GFP signal was much stronger than YFP, exposure time was set so that no YFP signal was detectable in images.
[f] *EXO70-VC* and *CDC24-VC* were constructed by the PCR-based C-terminal tagging method (Longtine *et al.,* 1998) using HP0038 pFA6a-VC-kanMX6 (Sung *et al.,* 2007) (a kind gift of W-K Huh, Seoul National University)

**Table S2. Plasmids used in this study**

| Plasmid | Description | Source |
|---|---|---|
| pHP2238 | pRS426-BEM1-YFP$^{N}$, 2μ, *URA3*, carrying the complete ORF of *BEM1* with 217bp upstream and 268bp downstream. | This Study |
| pHP2239 | pRS426-bem1$^{\Delta 1\text{-}147}$-YFP$^{N}$, 2μ, *URA3*, the same as pRS426-BEM1-YFP$^{N}$ except carrying a deletion of aa1-147. | This Study |
| pHP2240 | pRS426-bem1$^{\Delta 148\text{-}280}$-YFP$^{N}$, 2μ, *URA3*, the same as pRS426-BEM1-YFP$^{N}$ except carrying a deletion of aa148-280. | This Study |
| pHP2247 | pRS426-bem1$^{K482A}$-YFP$^{N}$, 2μ, *URA3*, the same as pRS426-BEM1-YFP$^{N}$ except carrying the mutation K482A in the PB1 domain. | This Study |
| pHP2248 | pRS426-bem1$^{\Delta 412\text{-}551}$-YFP$^{N}$, 2μ, *URA3*, the same as pRS426-BEM1-YFP$^{N}$ except carrying a deletion of aa412-551. | This Study |
| pHP2253 | pRS306-BEM1-YFP$^{N}$, integrative, *URA3*, carrying the complete ORF of *BEM1* with 217bp upstream and 268bp downstream. | This Study |
| pHP2254 | pRS306-bem1$^{\Delta 281\text{-}345}$-YFP$^{N}$, integrative, *URA3*, the same as pRS306-BEM1-YFP$^{N}$ except carrying a deletion of aa281-345. | This Study |
| pHP2255 | pRS306-bem1$^{\Delta 345\text{-}408}$-YFP$^{N}$, integrative, *URA3*, the same as pRS306-BEM1-YFP$^{N}$ except carrying a deletion of aa345-408. | This Study |
| pHP744 | YEp13, 2μ, *LEU2* | Broach *et al*., 1979 |
| PB268 | YEp13-RSR1$^{K16N}$, 2μ, *LEU2*, carrying the complete ORF of *RSR1* with the K16N mutation | Ruggieri *et al.,* 1992 |
| HP0038 | pFA6a-VC-kanMX6 | Sung *et al.,* 2007 |
